# Risk Factors Associated with Dengue Virus Infection in Guangdong Province: a Community-based Case-control Study

**DOI:** 10.1101/434472

**Authors:** Jundi Liu, Xiaolu Tian, Yu Deng, Zhicheng Du, Tianzhu Liang, Yuantao Hao, Dingmei Zhang

**Author notes:** Corresponding author: Dingmei Zhang, School of Public Health, Sun Yat-sen University, 74 Zhongshan Road II, Guangzhou, Guangdong, 510080, China.

## Abstract

**Background:** Dengue fever is a mosquito-borne infectious disease, and it is now still epidemic in China, especially in Guangdong Province. Owing to the absence of dengue vaccination, effective preventive measure is critical for controlling of dengue fever. This study aimed to explore the individual risk factors of dengue virus infection in Guangdong Province, as well as to provide a scientific basis for prevention and supervision of dengue fever in future.

**Methods:** A case-control study including 237 cases and 237 controls was performed. The data was collected from the epidemiological questionnaires. Univariate analysis was used for preliminary screening of 28 variables potentially related to dengue virus infection, and an unconditioned logistic regression analysis was used for multivariate analysis to analysis those statistically significant variables.

**Results:** Multivariate analysis of the result showed three independent risk factors: activities in the park (odd ratio [OR]= 1.70, 95%CI 1.03 to 2.83), outdoor sports (OR= 1.67, 95%CI 1.07 to 2.62), and poor indoor daylight quality (OR= 2.27, 95%CI 1.00 to 5.15); and two protective factors: two persons per room (OR=0.43, 95%CI 0.28 to 0.67), three persons and above per room (OR=0.43, 95%CI 0.22 to 0.86), using air-condition (OR=0.43, 95%CI 0.20 to 0.93).

**Conclusion:** These results are conducive to learn the risk factors for dengue virus infection in Guangdong Province. It is crucial to provide effective and efficient strategy to improve environmental protection and anti-mosquito measures. In addition, more systematic studies are needed to explore the other potential risk factors for dengue fever infection.

**Author summary:** Dengue fever, one of the mosquito-borne infectious diseases, is mainly transmitted by Aedes aegypti in Asia and Southeast Asia countries. Since 1978, the incidence of dengue fever has markedly increased in China especially in Guangdong province. In order to formulate the effective prevention and control measures, we explored the risk factors of dengue virus infection in Guangdong Province by conducting a case-control study. In this study, 237 patients with dengue virus infection and 237 participants without dengue virus infection were included. Then through these questionnaires and data analysis, we found that activities in the park, outdoor sports, and poor indoor daylight quality significantly contributed to the residents’ risk of dengue virus infection. On the other hand, we observed that using air-condition and using anti-mosquito measures were effective personal prevention interventions.

## Introduction

Dengue fever (DF) is an acute viral disease caused by four distinct serotype dengue virus, transmitted between humans by Aedes aegypti. In endemic countries in Asia and America, the burden of dengue is approximately 1,300 disability-adjusted life years (DALYs) per million populations, which is comparable to the disease burden of other childhood and tropical diseases, including tuberculosis, in these regions [1]. In Asia, epidemic dengue haemorrhagic fever (DHF) has expanded geographically from southeast Asian countries west to India, Sri Lanka, the Maldives, and Pakistan and then east to China [1]. According to the World Health Organization (WHO) statistics, there were only nine countries experiencing severe dengue epidemics before 1970. However, the number of countries which have experienced the disease now is more than 100, and the actual number of dengue infection is approximately 390 million, of which 500, 000 people require hospital admission because of severe dengue [2].

In 1978, the first reported DF outbreak due to Dengue virus (DENV) type 4 occurred in Foshan (adjacent to Guangzhou), a city in Guangdong Province, from where the spread of DF started in the southern provinces of China [3]. The DF incidence in Guangdong Province has been the highest in mainland China during the past decades, accounting for more than 65% of all cases in the country [4]. Besides, in 2013, total 4,662 cases occurred in mainland China, 62.08% of which were in Guangdong province. And the number of dengue fever cases increased dramatically to 48,162 in mainland China in 2014, 93.83% of these cases were reported from Guangdong province [5]. Thus, it is urgent to control the outbreak of dengue fever in Guangdong, which can serve as a bridge to transmit DF to other provinces in mainland China. Unfortunately, there are no effective drugs and vaccine to treat or prevent dengue fever so far.

In order to effectively prevent dengue fever, understanding the infection risk factors for dengue fever is necessary. Now, it is clear that the rapid growth of population, urbanization, and convenient modern transportation have greatly increased the spread of dengue fever [6]. However, most of the current case-control studies on risk factors are associated with sever dengue such as dengue shock syndrome (DSS) and dengue hemorrhagic fever (DHF) and the variables are all related to clinical and laboratory indexes [7–10]. In addition, with the development of the primary healthcare, some researchers were interested in exploring the positive effect of community participation in diminishing favorable household environments of dengue vectors [11,12]. Besides, environmental factors, awareness and knowledge of dengue prevention were also responsible for a significant reduction in dengue transmission [13,14]. To explore the dengue infection risk factors in Guangdong and provide basis for formulating control strategies in Guangdong province, several macroscopically descriptive studies were carried out, which provided more information on group level as well as climate factors, but less information on personal protective measures [15,16]. In order to get more related information of risk factors on individual level and provide more specific prevention approaches, this case-control study was conducted. Geographically, Guangzhou city and Zhongshan city are located in the Pearl River Delta Region of Guangdong, which is the main area where dengue fever epidemics highly [17,18], meanwhile, Guangzhou city is the capital of Guangdong province and the first reported case of autochthonous dengue fever occurred in Zhongshan city [19]. Thus, the prevalence of dengue fever in Guangzhou city and Zhongshan city is a good representation of Guangdong’s. The cases and controls, which were selected from the communities in Guangzhou city and Zhongshan city, were identified based on serum test of dengue virus IgG and IgM. The potential risk factors analyzed included personal life activities, personal hygiene habits, housing situation, living environment, mosquito protection status and residential surroundings.

## Methods

### Samples collection

The case-control study was performed based on the project of Research on Prevention and Control of Human Immunodeficiency Virus and Hepatitis B Virus in Guangdong province which has a 200,000-sample database. The demographic information of the database could be seen in our related study [20]. Totally, 3,136 serum samples were selected from residents of Yuexiu District in Guangzhou city (699), Liwan District in Guangzhou city (1386), Torch Development Area in Zhongshan City (180) and Xiaolan Town in Zhongshan city (1051) respectively via stratified cluster sampling rooted in the database for serological testing. That is, approximate 30 to 35 persons a month were sampled from every age group (under 19 years group, 19 to 40 years group, 41 to 65 years group, and over 65years group) during two years from September 2013 to August 2015.

### Enzyme Immunoassay Test

The enzyme-linked immunosorbent assay (ELISA) was used to detect dengue antibody IgG and IgM. The IgG antibody was measured by indirect ELISA (LOT: 01P20A006, Inverness Medical/Panbio Australia), whereas the antibody of dengue IgM was tested via capture ELISA (LOT:01P30A002, Inverness Medical/Panbio Australia). The undefined results were confirmed by the colloidal gold method (LOT: DEN141001, Inverness Medical/Cortez USA). The detail could be found in the relevant study [20].

### Ethical Statement

The work obtained the ethics approval from the Institutional Review Board of the School of Public Health at the Sun Yat-sen University (L2017030), in line with the guidelines to protect human subjects. After understanding the research objective and being ensured that their personal information can keep private, all research subjects or the guardians signed the written informed consents. And every of the participants had the right to withdrawal from this study at any time.

### Case definition and selection of control

Among the 3,136 serum samples, the number of individuals detected as IgG antibody positive and IgM antibody positive were 305 and 103 respectively, 34 of which were positive for both antibodies. Thus 374 antibody-positive samples were defined as dengue infection and those willing to receive questionnaire survey were chosen as members of the case group.

In this study, 256 dengue infections were chosen to fill in the questionnaires, however, 19 questionnaires with missing most of important information were eliminated. Eventually, 237 infections were included in the case group. There was no statistical difference in gender between the persons who were willing to receive questionnaire survey and those who were unwilling (*p*=0.950>0.05), as well as in age (*p*=0.127>0.05).

The controls were selected from those who were tested negative both to IgG and IgM using frequency matching. That is, the candidate controls were stratified according to the age and sex ratio of the case group, and selected by convenience sampling (participants volunteered to be part of the samples) from each layer. In total, 308 questionnaires were completed by the persons whose IgG and IgM antibodies were both negative. To enhance the comparability of the study, according to the community information of the 237 cases, a further match was done and 237 controls were selected.

### Data collection and analysis

The phone questionnaire was carried out by these trained investigators to obtain the information strictly according to the facts, then some subjects were interviewed face to face to verify the validity of their information. The main contents of the questionnaire included: general demographic characteristics (age, gender, blood type, average household income); personal life activities, such as activities in the park, outdoor sports (such as hiking, mountain climbing and camping) and outbound tourism experience; personal hygiene habits (domestic sewage and garbage management, participate in the community hygiene management intervention); housing situation, like the age and area of inhabitation and living floor; living condition (average numbers of person per room, use of air conditioning, indoor daylight quality, animals or aquatic plants on property, etc.); mosquito protection status (use of mosquito nets, pesticide, etc.); residential surroundings (whether there are junk yards, ponds or construction sites within 200 meters).

Epidata3.1 software was used to establish a database of individual risk factors for dengue infection among residents in Guangdong Province. All the data were analyzed by SPSS statistics 23.0 software. A univariate analysis was applied for preliminary screening of variables, and there would be an unconditioned logistic regression analysis method for multivariate analysis to analysis those statistically significant variables.

## Results

### General demographic characteristics of the samples

In total, 474 subjects were recruited successfully, including 237 cases and 237 controls. The gender ratio was 1:1.66 (male: female) in both the case group and the control group. The potential confounders were comparable between the two groups (Table 1).

**Table 1.**
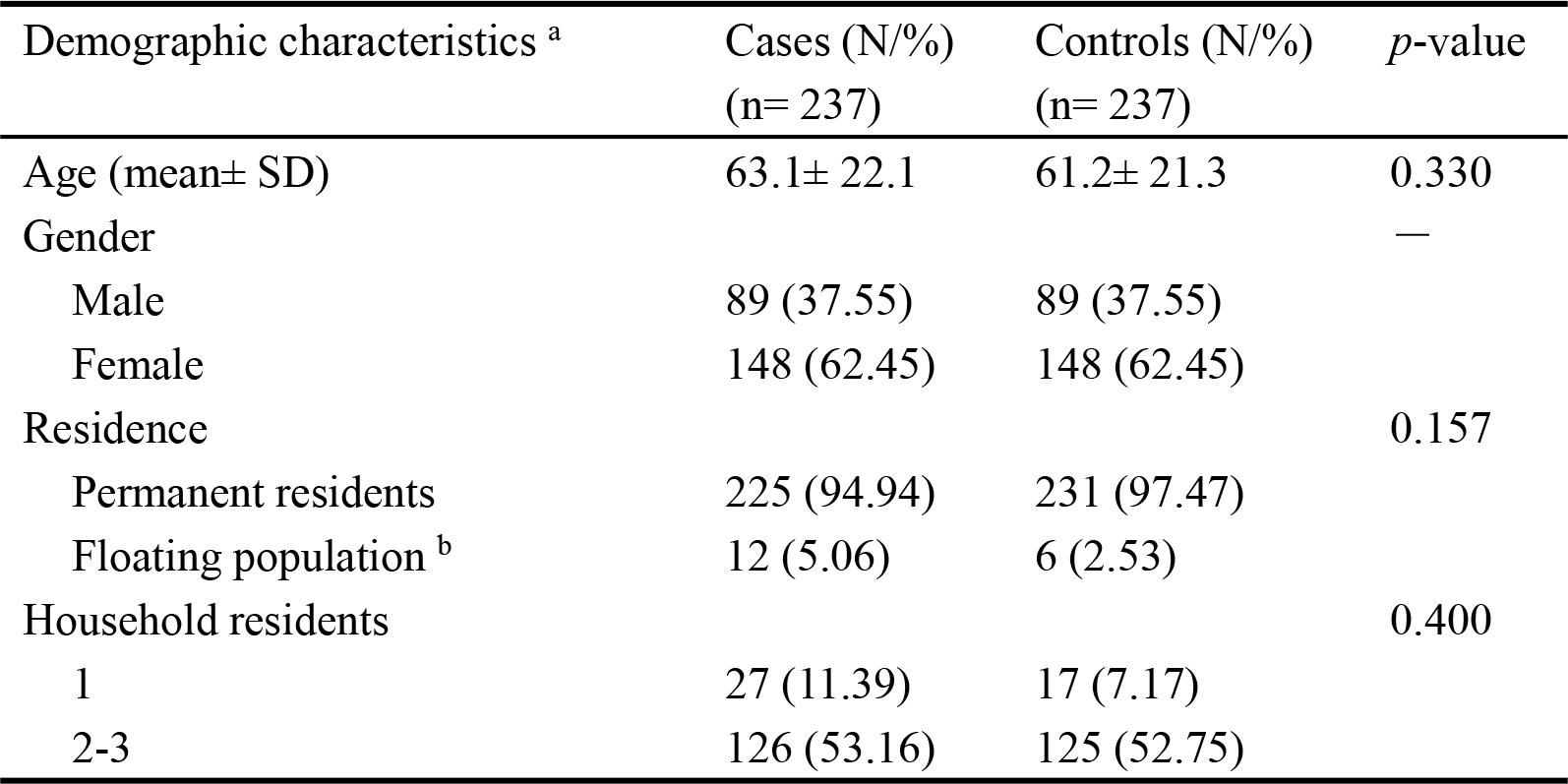
The demographic characteristics of cases and controls

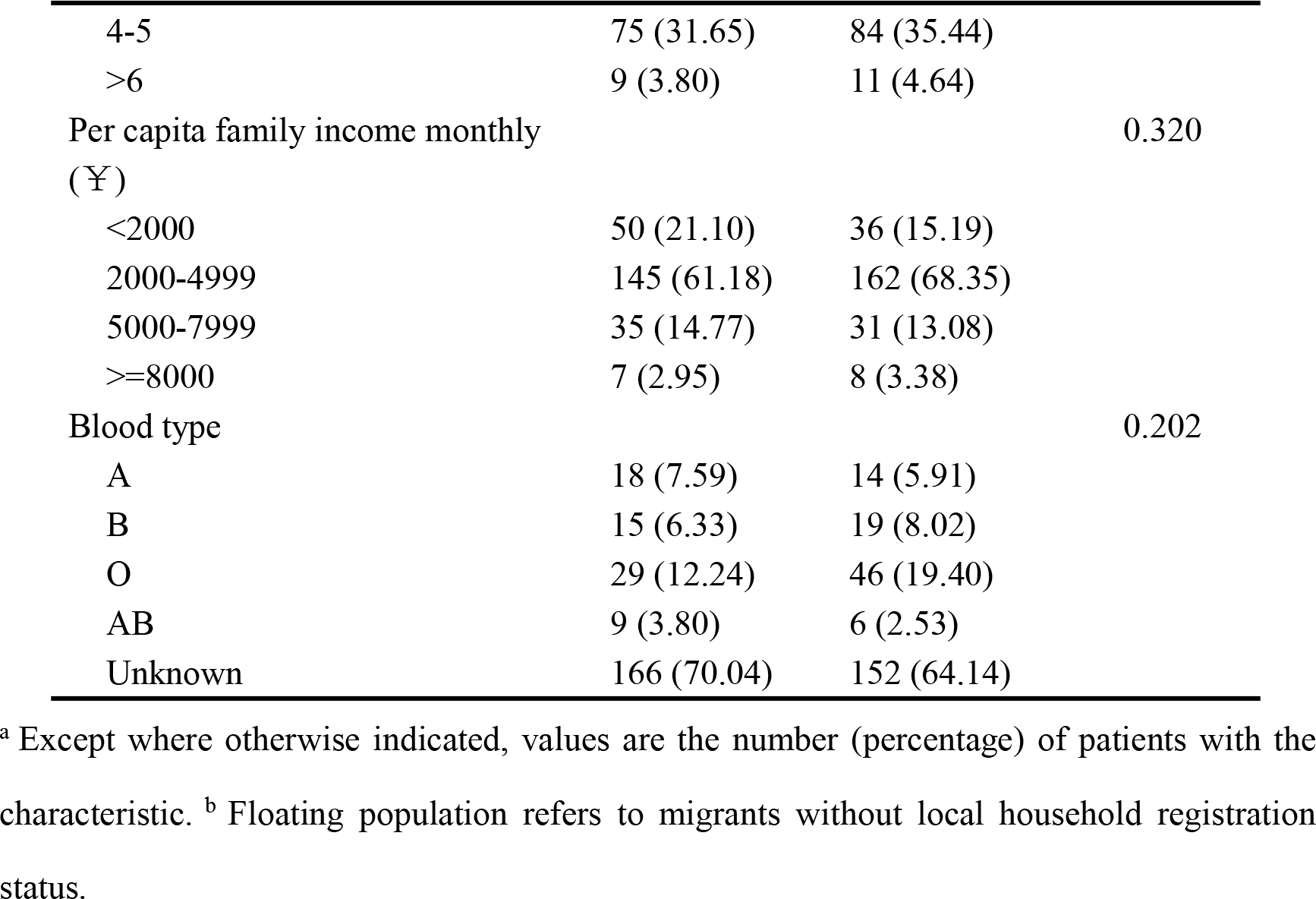

### Univariate analysis

Total 28 potential risk factors were analyzed, which were further divided into six dimensions, which were personal life activities, personal hygiene habits, housing situation, living conditions, mosquito protection status and residential surroundings respectively. Just as shown in Table2, people who with activities in the park had a significantly higher risk of getting dengue virus infection than those who without activities in the park (*p*=0.049<0.05). Significantly, people who had outdoor sports were more likely to be infected by the dengue fever virus compared with those who had no outdoor sports (*p*=0.009<0.05). At the same time, there were statistical difference on housing type (*p*=0.040<0.05), the average numbers of person per room (*p*=0.000<0.05), using air-condition (*p*=0.026<0.05) and the indoor daylight quality (*p*=0.032<0.05) between the case group and the control group.

But rather, for the factors of residential surroundings within 200 meters, there were no statistical difference on the existence of garbage collection sites, junk yards, ponds, and construction sites.

**Table 2.**
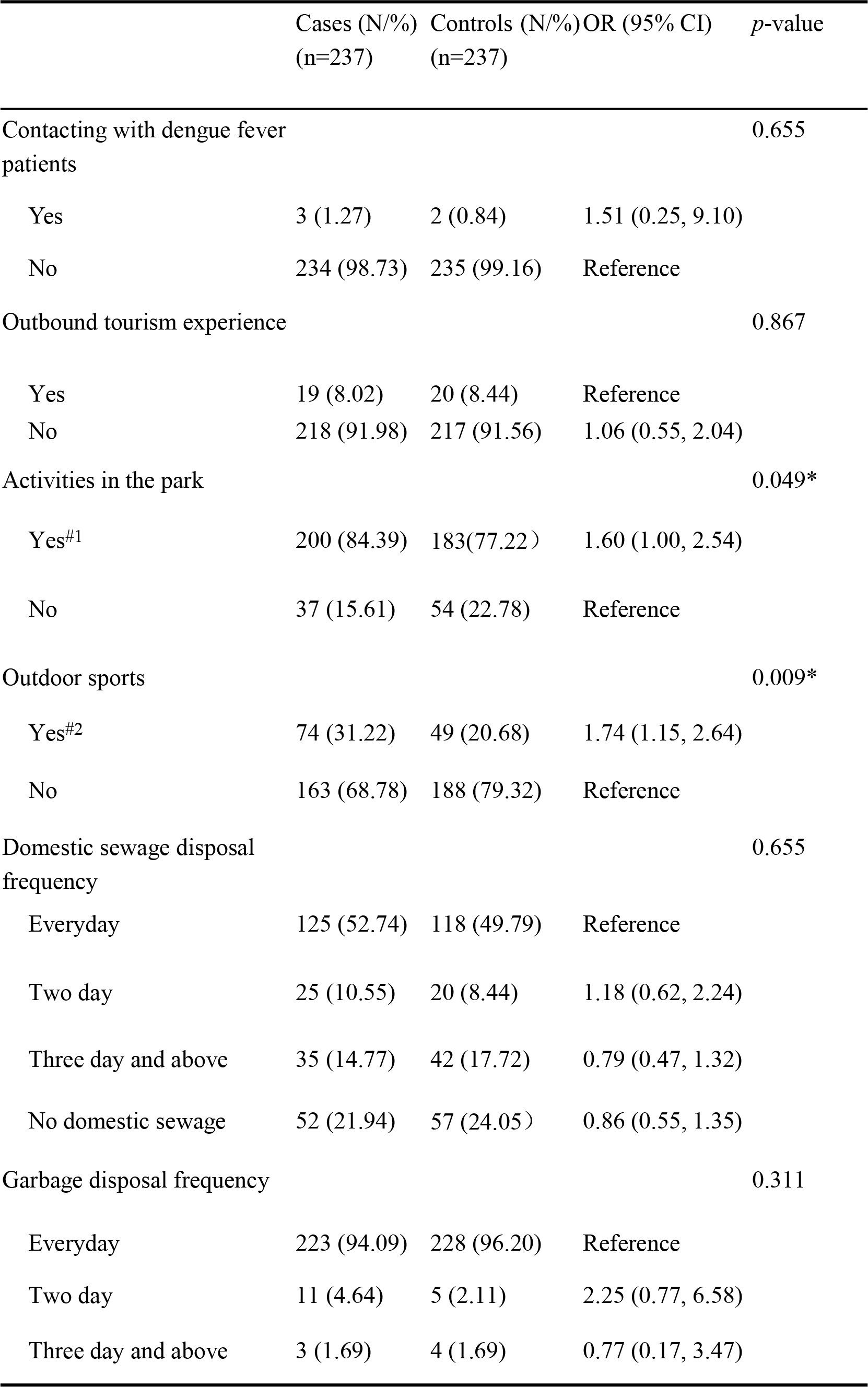
Univariate analysis of risk factors for dengue virus infection

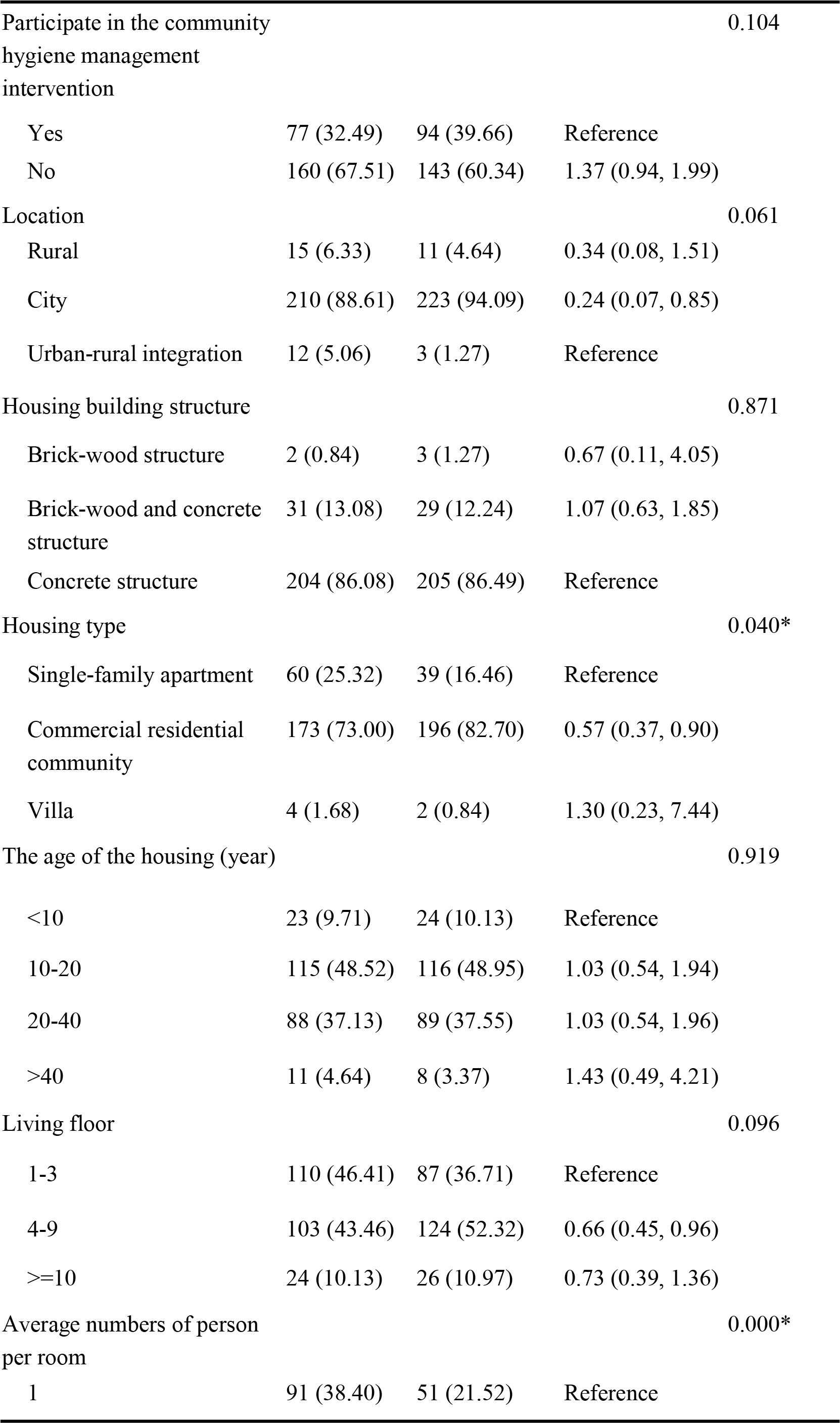

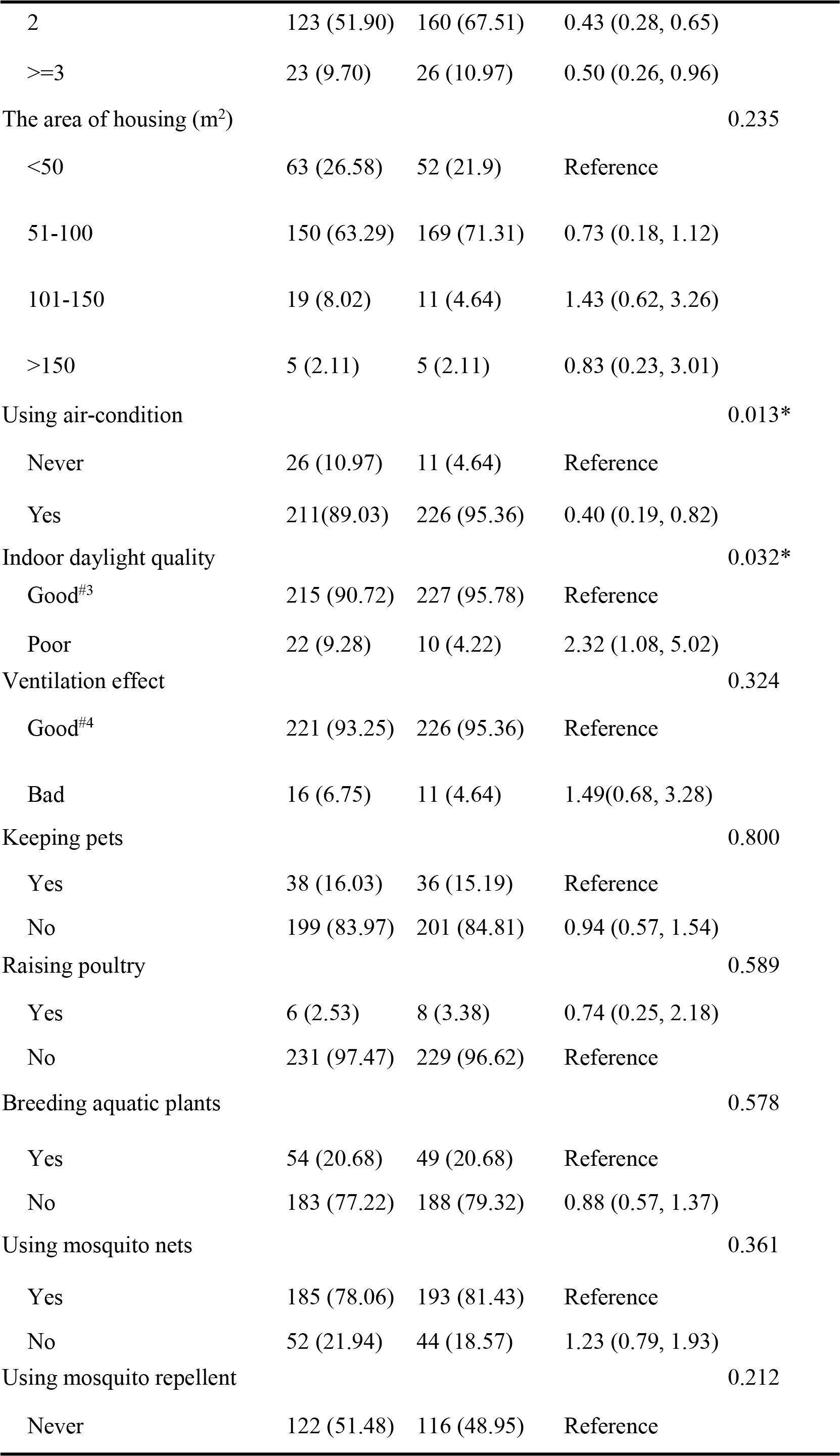

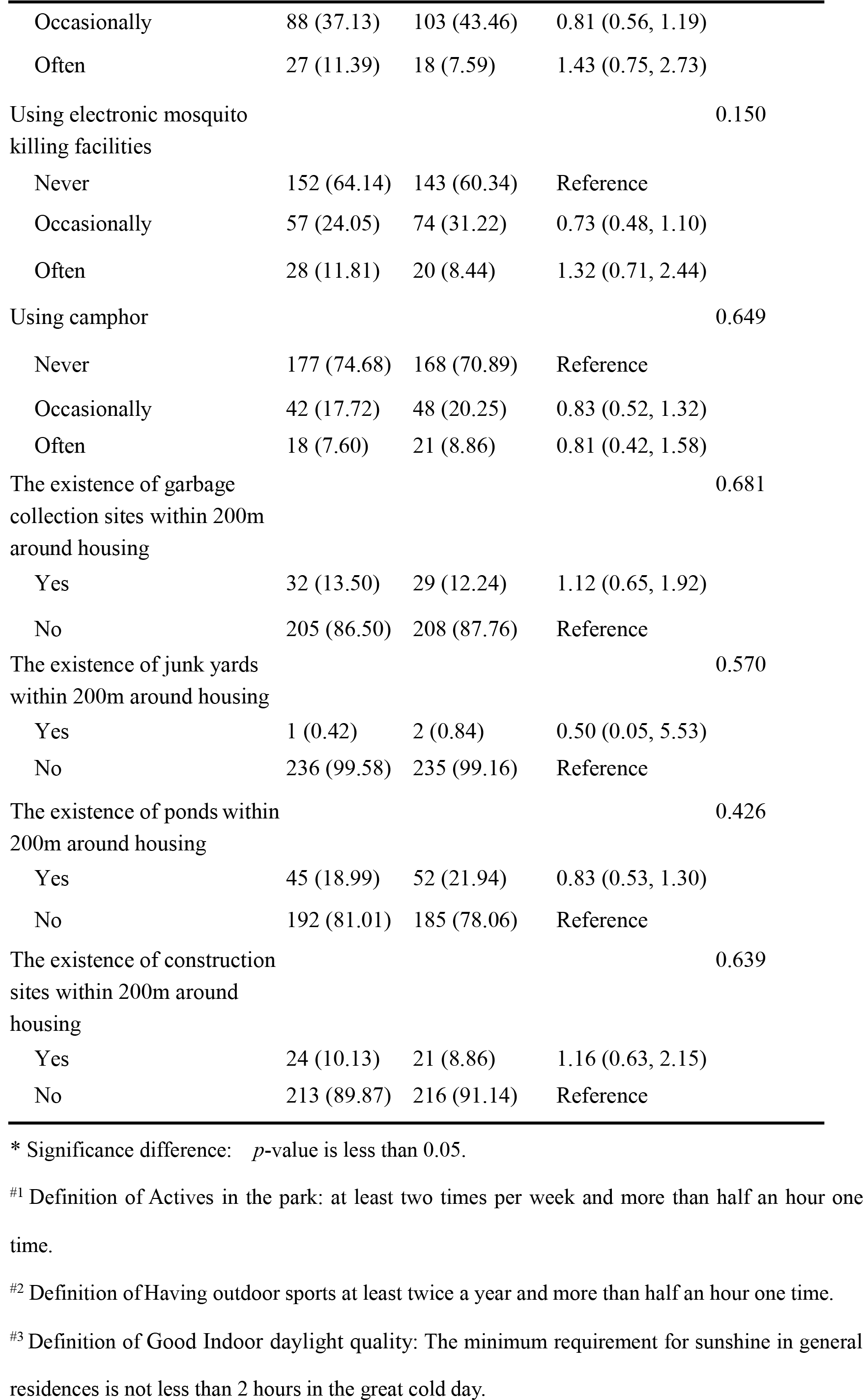

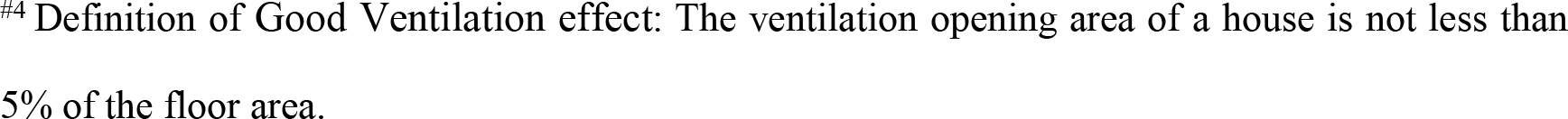

### Multivariate analysis

Through the univariate analysis above, there were statistical significance on six variables between the case group and the control group (*p* values are all less than 0.05). To further analyze the relative importance of the six various factors associated with dengue infection, they were all brought into a multivariate model. In the unconditioned logistic regression model, activities in the park, outdoor sports and the poor indoor daylight quality were significantly associated with increasing risk of dengue virus infection, with odd ratios (ORs) of 1.70, 1.67, and 2.27 respectively. On the other hand, two persons per room (OR=0.43), three persons and above per room (OR=0.43) and using air-condition occasionally (OR=0.43) were significantly associated with protections against dengue virus infection (Table 3). The equation obtained by unconditional logistic regression is: ln (Odds) = 1.193 + 0.533 X_1_ + 0.515 X_2_ − 0.835 X_3_^a^ − 0.835 X_3_^b^ − 0.837 X_5_ + 0.822 X_6_. X_1_ presents park experience, X_2_ presents outdoor sports, X_3_^a^ presents two persons per room, X_3_^b^ presents three persons and above per room, X_5_ presents using air-condition and X_6_ presents the poor indoor daylight quality.

**Table 3.**
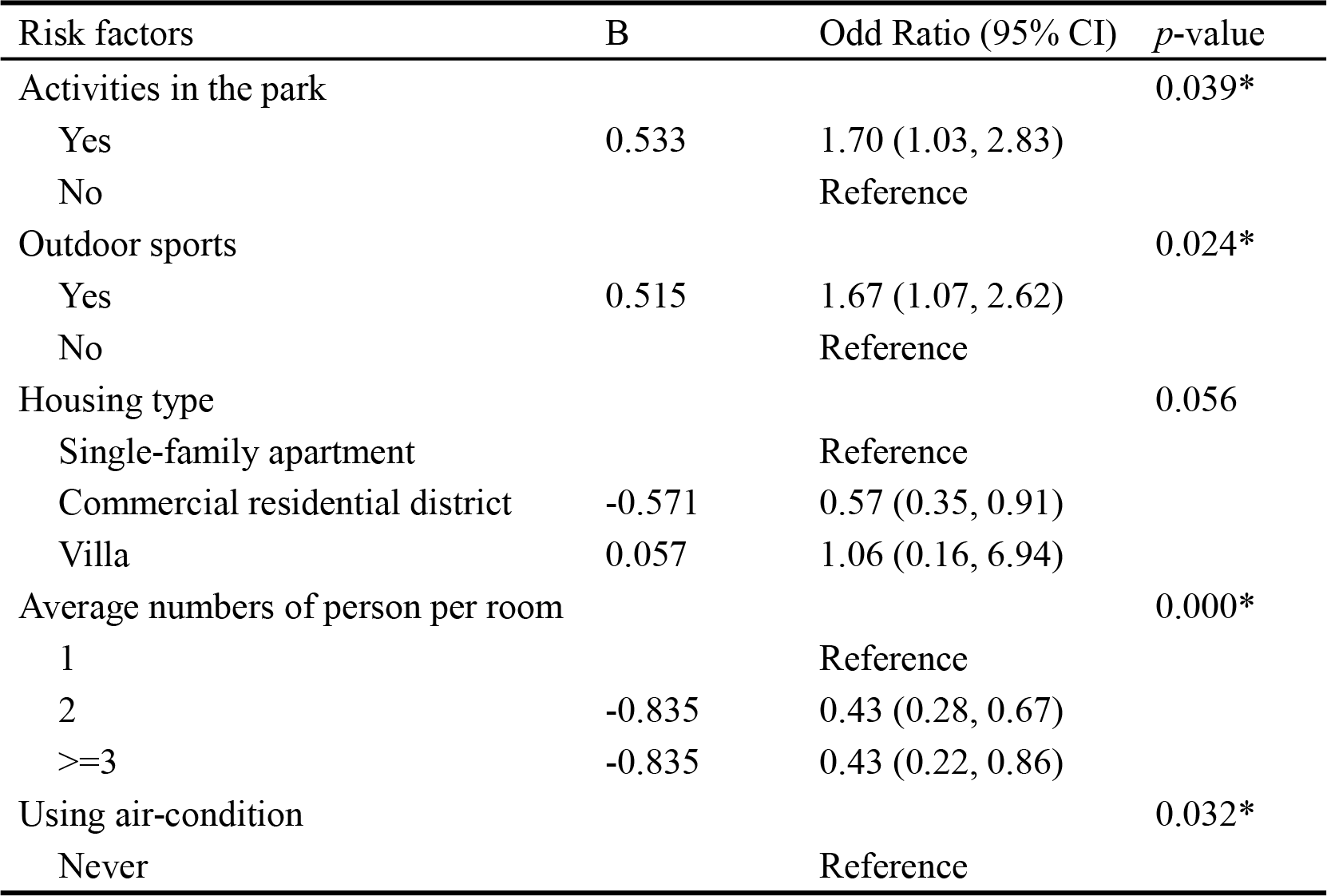
Multivariate analysis on risk factors for dengue virus infection

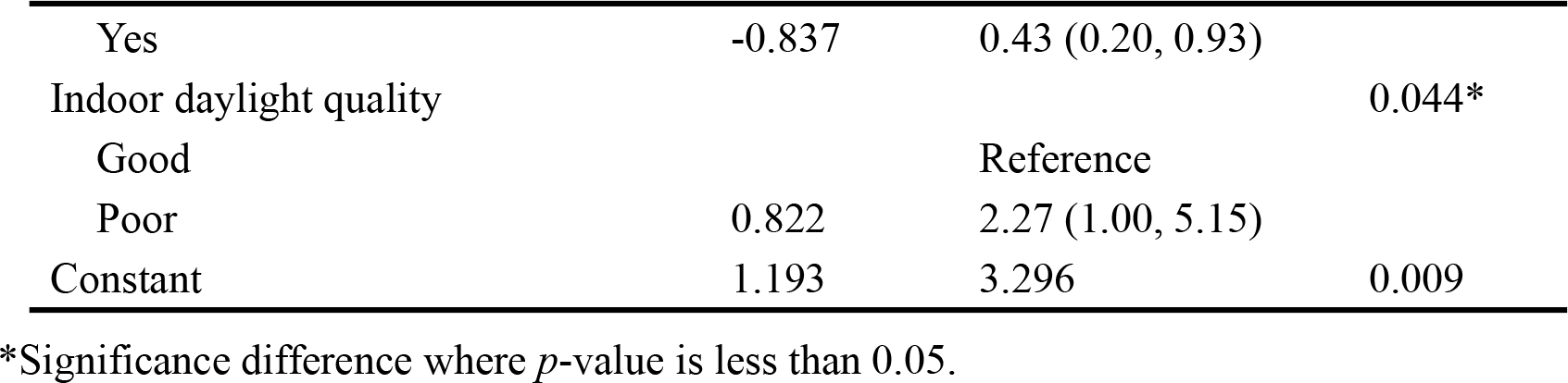

## Discussion

Our study showed that activities in the park, outdoor sports and the poor indoor daylight quality significantly increased the 1.70-time, 1.67-time, and 2.27-time risk of developing dengue infection respectively in Guangdong Province. In addition, it also suggested that more than one person per room and using air-condition might decrease the risk for dengue virus infection by close to 0.43 times.

The result of the study revealed that people with activities in the park have significantly higher risk of contracting dengue fever than those with no activities in the park. This could be related to the high density of the mosquito of Aedes albopictus in the park [21]. And when playing in the park especially in the morning and at dusk, residents were usually casually dressed with more skin exposed, which made it easier to be bitten by mosquitoes and increased the risk of dengue infection. Besides, research showed that the main protocols of mosquito prevention were using mosquito nets, pesticide and mosquito repellent, but less residents tend to use mosquito repellent outdoor [22]. Thus it is necessary for residents to increase the awareness of adopting some approaches to prevent mosquito in their daily lives and reduce the risk of dengue fever. For instance, wearing long-sleeve clothes during activities in the park. Meanwhile, having outdoor sports was another risk factor for dengue virus infection, the explanation for which might be that the forest margin, the holes of trees and the natural reservoirs were the origins of Aedes albopictus. Simultaneously, these places were good choices for residents to hike and go camping [23]. Therefore, outdoor sports increased the risk of mosquito bite, and it was necessary to adopt anti-mosquito measures such as insecticide treated materials in the outdoor.

We also found that the density of human population was closely associated with dengue transmission. In general, it is believed that the high population density is a risk factor for dengue transmission [6,24]. But in our study, those sharing a crowded household with 2 persons and 3 persons and above were less likely to have a dengue infection. The conclusion, however, was contrary to Velascosalas ZI, et al.’s research [25]. The explanation could be that most of one room with 3 persons and above were shared by the parents and their young children which was related to Chinese way to raise kids. When the parents with their kids lived in one room, they would pay more attention on using anti-mosquito measures and maintaining a good sanitary environment to avoid the kids being bitten. Secondly, in our related study, the result showed that the married group had a lower rate of infection than the widowed group and the divorced group [20], it also suggested that the married group who usually living in one room with 2 persons and above had a less risk to be infected by dengue virus. Thirdly, when the number of mosquitoes was fixed, the more persons in one room, the lower probability of a person being bitten by the mosquito. Therefore, further research is needed to determine whether more than one person per room is a protective factor or a risk factor for dengue virus infection.

According to Shen et al. [26] and Wu PC et al.’s [27] researches, the yearly average temperature higher than 18°C would increase the risk of dengue virus infection. Meanwhile, our study indicated that using air-condition was a protective factor against dengue infection. This reason might be that using air-condition to cool indoor environment can reduce the risk of dengue transmission. Secondly, when using air-condition, doors and windows were commonly shut down, which reduce the chance of mosquitoes entering the room.

Besides, our study showed that the poor quality of indoor daylight increased a 2.27-fold risk for dengue virus infection. The explanation for this is possibly that adult Aedes albopictus prefer to inhabit in weak-light places rather than in those areas with plenty of light [28,29]. So the environment of poor daylight was suitable for the survival of mosquitoes, which lead to the high density of mosquitoes in the room.

Our result failed to find the protective effect of mosquito bed nets on reducing the risk of dengue fever infection. And the result was consistent with Tsuzuki A et al. [30] and Loroñopino MA et al.’s [31] studies. One reason might be that these mosquito bed nets were usually used at night. However, the mosquito of Aedes albopictus was found biting throughout daylight hours, especially in the early morning and late afternoon [32]. Another explanation could be that the majority of residents including the case group and the control group chose to use mosquito bed nets to prevent mosquito bites, which might lead to lose statistical significance between using mosquito bed net and dengue infection. The third cause might be that good living environment, for instance, the popularity of air-condition and mosquito killing facilities, reduced the demand for mosquito bed nets. Despite the limitation, based on our local experience and some studies’ results [33,34], bed nets are still recommended to use not only at night but also during the day [22].

Andersson N et al. [35] and Roberto TC et al.’s [12] studies revealed that the government’s vector controlling capacity had a very important impact on dengue transmission. Because of the high incidence of dengue fever in Guangdong province in recent years, the community neighborhood committees and property management departments organized many health remediation activities under the supervision of the relevant health agent or the Centers for Disease Control and Prevention (CDC) [36], greatly improved the residential living environment, reducing mosquito breeding, which may be an explanation of that the variables of domestic sewage disposal, garbage management and residential surroundings were not statistically significant in this study. However, 67.51% of the infections and 60.39% of the controls didn’t participate in the community hygiene management intervention activities in neighborhood committees or property management organizations in this study. Regardless of the case group or the control group, their public health consciousness need to be strengthened. If the government has a good macro-control system without the support of the masses, it will be unable to fight against disease and establish a sound prevention system.

## Limitations

There are several flaws in this study that should be overcame in the future study. First, cases and controls were identified according to the result of antibody detection. However, the IgG-positive samples might be infected a few years ago, which led to the results of their completed questionnaires were incompatible with the situations when they were infected. Besides, the existence of recall biases which was caused by the inaccuracy of memory could also reduce the authenticity of the questionnaire. Second, in an initial infection, the titers of the IgG antibody was so low after having a fever that the antibody couldn’t be detectable [37], thus the newly infected persons was misdiagnosed as the members of the control group. As a result, the misclassification bias was brought. To reduce these bias, some of the questionnaires were completed by the community doctors, and some of the questionnaires were completed by face-to-face interviews. Third, the data was obtained from serum sample of residents in Guangdong Province from 2013 to 2015 without other extrapolated studies, the model’s general practical application is difficult to evaluate at present.

The focus of our study is on residents in communities with mild or asymptomatic dengue virus infection, rather than patients with severe clinical symptoms, which is more representative to explore the risk factors for dengue virus infection in Guangdong province. Besides, this study shows the relationship between dengue infection and individual risk factors, which was beneficial for avoiding being infected by dengue virus. Meanwhile, the results of this study provided clue and basis for dengue fever prevention and control.

## Conclusion

The case-control study revealed some risk factors for dengue virus infection to provide guidance for the concrete preventive measures. After analyzing the 28 relevant variables, it is easy to find that when having the behaviors of activities in the park and outdoor sports, it is useful to use mosquito control measures and reduce the exposed area of skin to low down the risk of dengue infection. Moreover, improving the quality of indoor lighting and using air-condition can get the same effect. In the long run, more variables need to be introduced into the logistic regression model and further research should be conducted to provide theoretical basis for formulating prevention and control measures of dengue fever.

## Acknowledgments

We are grateful to the National Science and Technique Major Project for offering assistance in sample collection; the research assistance is offered from the Health and Family Planning Commission in Yuexiu District and Liwan District, Yuexiu District Centre for Disease Control and Zhongshan Centre for Disease Control; the investigators such as staff and students in the School of Public Health for providing field and laboratory assistance, and all subjects.

## Supporting information

S1 Dataset. Complete data.

S1 Checklist. Strobe checklist - case control.

